# Low-cost and highly efficient generation of near-complete bacterial pathogen genomes by TELL-Seq

**DOI:** 10.1101/2024.08.17.608388

**Authors:** Zhuoying Wu, Ting Zhang, Sheng Li, Lingzhu Shen, Zhangnv Yang, Yunfeng Yang, Xuanheng Chen, Bolin Li, Shicong Zhou, Xintian Zhou, Beibei Wu, Jianmin Jiang, Xiao Li

**Author notes:** **Corresponding author:** Beibei Wu, Jianmin Jiang, Xiao Li. These authors contributed equally to this work (Zhuoying Wu, Ting Zhang).

## Abstract

Recent evidences suggest that *de novo* genome assembly can provide additional insights into genomic variation landscape beyond short-read NGS analyses. Despite the advent of longread sequencing technologies, generating high-quality bacterial genome assemblies remains expensive and requires high-quality and large quantities of DNA. TELL-Seq, an emerging linked-read technology, has been reported as a low-cost alternative for producing near-complete genome assembly in certain bacterial species. However, a systematic assessment of its performance and characteristics in *de novo* bacterial genome assembly has not yet been conducted. To address this, we benchmarked TELL-Seq using a set of clinical and standard bacterial pathogens with a wide range of genome size (2.0 - 6.4 Mbp), GC-content (32% - 68%), genome complexity (mappability from 0.15 to 0.60) and Gram types. Our findings indicate that, with 1/6 cost and 1/15,000 of DNA compared to PacBio HiFi, TELL-Seq could generate as high quality near-complete bacterial genome assemblies. The optimal depth for assembly is around 200×, as increased depth improves contiguity and completeness, though quality plateaus at 250×. In general, genome complexity was significantly correlated with assembly quality, but GC-content was not. Nevertheless, genomes with extreme GC-content may require higher depths for accurate assembly. Our study suggests that TELL-Seq could be a scalable method for large-scale bacterial genomic surveys. The data generated from this study could serve as a benchmark dataset for further algorithm development.

**Impact Statement:** An increasing body of literature have shown that *de novo* genome assembles could provide additional insights into genomic variation beyond standard short-read technologies. It’s particularly useful in clinical and epidemiological studies of bacteria, due to high variable nature of their genomes. Although great advances have been made in long-read sequencing technology, it’s still expensive with high requirement over sample DNA’s quality and quantity, limiting its adoption in large-scale studies or surveys. Linked-read technology is an inexpensive alternative which used single molecule barcoding to mark origin of reads for better scaffolding of contigs assembled from normal short reads. TELL-Seq, as one of these promising technologies, was reported to be able to inexpensively and efficiently generate near-complete bacterial *de novo* genomic assemblies with low DNA inputs. There’s not yet a systematic benchmarking of this technology in bacterial pathogens. In this study, a set of clinically important bacterial pathogens with a wide spectrum of GC-content, genome size, genome complexity, and Gram Type were used to investigate the performance of TELL-Seq technology in *de novo* genome assembly. We found that, with about 1/6 of cost and 1/15000 of DNA quantity, it could generate comparable *de novo* assemblies compared to standard reference genomes or PacBio HiFi assemblies, while beating conventional short-read assemblies. We found several key factors, such as genome complexity, and sequencing depth, which would impact the quality of the assembly. Our study suggests that a sequencing depth of 200× is sufficient to achieve satisfactory results. Our study has revealed characteristics of this technology in bacterial genome assembly and paved the way for its large-scale application in clinical or epidemiological surveys.

**Data Summary:**

1. Raw sequencing data for all isolates have been deposited to SRA, accessible through these NCBI BioProject PRJNA1133244.
2. A full list of SRA BioSample accession numbers are available in Tab S5.
3. Assemblies and analyses companion this paper is available at: https://github.com/x-lab/tellseq_bacteria

## Introduction

The advent of long-read sequencing technology has underscored the importance of *de novo* assemblies in genomic studies [1], allowing for the discovery of genomic variations that conventional NGS technology often misses. In eukaryotes, many studies have successfully assembled telomere-to-telomere genomes, which have facilitated the construction of pan-genome models for variant discovery [2-5]. While the concept of pan-genomes originated from characterizing bacterial core and accessory gene sets, it has expanded to include all genome sequences, represented by graph data structures [6].

In bacterial genomics, there is growing interest in studying nucleotide-resolution pan-genomes [7]. Despite the rapid advancement of long-read sequencing technologies, their high cost and demanding DNA quality and quantity requirements limit their application. Technologies capture long-range information using short sequencing reads have been developed in the past decade to overcome the weakness of long-read and short-read sequencing technologies. These include synthetic long reads (SLR) sequencing, such as Illumina TrueSeq [8], and linked-read technologies, like MGI’s stLFR [9], Element’s LoopSeq [10], Universal Sequencing Technology’s TELL-Seq [11], and 10× Genomics’s linked-reads technology [12]. Linked-read technologies enhance short-read NGS performance by linking all sequencing reads that come from the same long DNA fragment with unique barcode. This scaffolding information improves *de novo* assembly results of short NGS reads.

Linked-read sequencing performs comparably or even better than long-read sequencing in bacteria, which have simpler genomes compared to eukaryotes [13]. Near-complete bacterial *de novo* genome assembly has been achieved using TELL-Seq in some bacteria species [11,13-14]. As infectious diseases caused by bacterial pathogens remain a significant public health threat, there are growing advocacy for using pathogen genomics to ensure consistent and timely monitoring and control. Timely and cost-effective near-complete bacterial genomes enhance study accuracy, and linked-read technology shows great potential for this purpose. However, systematic evaluations and insights into key factors influencing *de novo* assembly quality are lacking.

To address this, we selected a diverse range of standard and clinical bacterial pathogens, varying in GC-content, genome size, genome complexity, and Gram type, to benchmark the performance of TELL-Seq. TELL-Seq effectively assembled near-complete genomes in most cases, but caution is advised for highly complex genomes. We also provide recommendations on suitable sequencing depths and highlight factors influencing assembly quality. Our dataset can support further algorithmic development of linked-reads assembly.

## Results

### Selection of Diverse Bacterial Pathogens for Systematic Benchmarking

To systematically assess the performance of TELL-Seq in *de novo* assembly of bacterial pathogen genomes, we selected 12 clinically relevant bacterial species, ensuring a broad range of GC-content (32% - 68%), genome sizes (2.0 Mbp - 6.4 Mbp), genome complexity (mean mappability from 0.15 to 0.60), and Gram types (50% Gram-positive) (Tab S1). For a comprehensive evaluation applicable to both research and clinical settings, we included one standard ATCC isolate and one corresponding clinical isolate for each species. One clinical sample did not meet quality control standards and was excluded from further analyses. We evaluated the quality of TELL-Seq-based *de novo* assemblies by comparing them against reference genomes for the standard isolates and PacBio HiFi-based assemblies for the clinical isolates.

### TELL-Seq produces near-complete genome assemblies at 1/6 cost of PacBio

To evaluate the quality and cost-effectiveness of TELL-Seq for studying bacterial pathogens, we compared *de novo* assemblies of clinical pathogens using Illumina, TELL-Seq, and PacBio technologies. Our analysis revealed that TELL-Seq produced near-complete assemblies comparable to those achieved by PacBio, with an average genome coverage of 97.4%. The average fragment length for TELL-Seq was 10,128 bp, closely approaching PacBio’s average read length of 13,601 bp, and the longest contig covered 95% of the reference genome. Notably, TELL-Seq achieved these results at only one-sixth of the cost ($107 vs. $682) and required just 1/15,000th of the DNA input (Tab 1). While Illumina short-read sequencing was the most cost-effective at $62 per sample, its shorter read length led to more fragmented assemblies, compromising assembly contiguity (Tab 1).

**Table 1:** Sequencing and assembly quality for bacterial pathogens using different technologies.

|  | Illumina | TELL-Seq | PacBio |
| --- | --- | --- | --- |
| gDNA input (ng) | 0.1 | 0.1 | 1500 |
| Total base (Gbp) | 33.91 | 33.91 | 25 |
| Average base ± SD (Gbp) | 3.1 ± 0.2 | 3.1 ± 0.2 | 2.3 ± 0.97 |
| Average read length* ± SD (bp) | 119 ± 3 | 10,128 ± 2,733 | 13,601 ± 2,672 |
| Average genome fraction (%) for 1 longest contig | 17.4 | 95.0 | 1 |
| Average genome fraction (%) for 2 longest contigs | 29.3 | 95.7 | 1 |
| Average genome fraction (%) for 3 longest contigs | 37.4 | 96.0 | 1 |
| Average genome fraction (%) for 4 longest contigs | 44.0 | 96.0 | 1 |
| Average genome fraction (%) for 5 longest contigs | 49.1 | 96.2 | 1 |
| Average genome fraction (%) for all contigs | 97.5 | 97.4 | 1 |
| Average N50/reference length | 0.15 | 0.96 | 1 |
| Average contig number (≥ 0 bp) | 115 | 61 | 2 |
| Average contig number (≥ 1000 bp) | 65 | 30 | 2 |
| Average contig number (≥ 5000 bp) | 49 | 4 | 2 |
| Average contig number (≥ 10000 bp) | 37 | 3 | 2 |
| Average contig number (≥ 25000 bp) | 23 | 3 | 2 |
| Average contig number (≥ 50000 bp) | 14 | 2 | 1 |
| Average Cost per sample (US\$) | 62 | 107 | 682 |

Read length\*: For TELL-Seq, read length is size of Super Long Fragment (SLF)

### Genome complexity, genome size and sequencing depth are the key factors influencing TELL-Seq genome assembly quality

We further went on to identify factors that are crucial for the quality of TELL-Seq *de novo* assemblies. To begin with, we accessed the impact of sequencing depth, one of the most fundamental and cost relevant factors, to assembly quality. Through a down-sampling experiment, we noticed that there was a clear positive correlation between sequencing depth and assembly quality (Fig.1b), indicating that sufficient depth of coverage is critical to *de novo* assembly. However, this trend was non-linear, with a clear plateau effect around 200×. At 250× depth, a slight decline in assembly quality was observed. This might be due to limited ability of the Tell-Link assembly algorithm in handling over-deeply sequenced reads.

We also noticed that at same depth of coverage, some genomes, such as *N. meningitidis*, had inferior assembly results compared to other species (Fig. 1a). We hypothesized that other factors, such as genome complexity, genome size and GC-content, might also influence the genome assembly quality. To further look into this, we asked if there were significant correlation between genomic characteristics (i.e. mappability, GC-content and genome size) and assemble quality (contig number, genome fraction, and N50/genome length). Minimum mappability displayed a significantly high correlation with assemble quality at almost all depth, with absolute correlation coefficient >0.5 and p-value < 0.05. As the sequencing depth increased, the assembly quality became less dependent on the genome complexity, with an inflection point observed at 200× for both contig number and the N50/genome length ratio (Fig. 1b, Fig. S3, Tab S4). This finding suggests that while an increase in sequencing depth may generally enhance assembly quality, in most cases, a depth of 200× was sufficient to achieve optimal correlation with the assessed assembly metrics. Some extent of correlation was found between genome size and assembly quality (Fig.1b), ascertained with Gram-type (Fig.S3). We did not observe any significant correlation between GC-content and assemble quality (Fig. 1b, Fig. S2, Tab S4).

**Fig 1:**
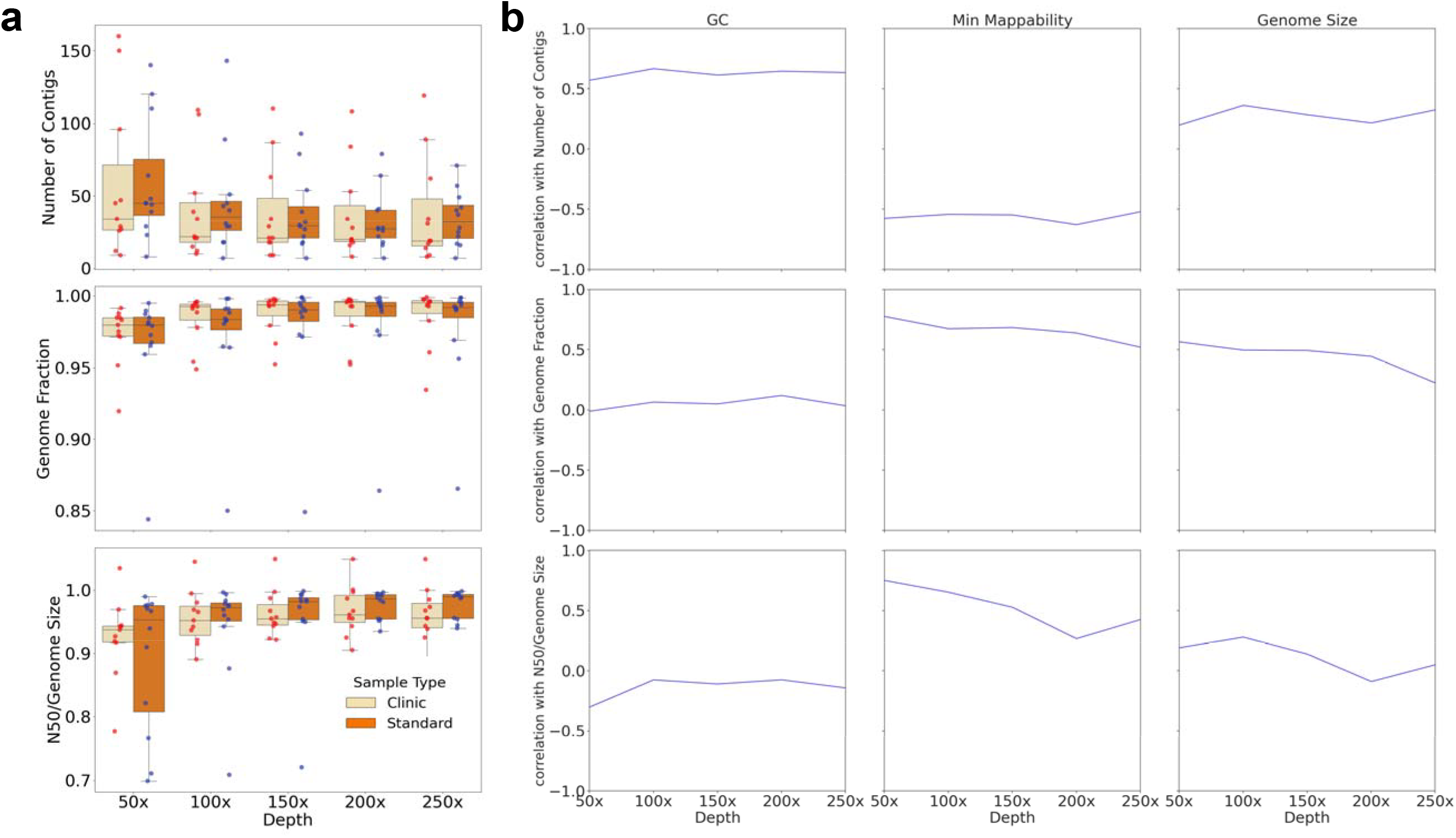
TELL-Seq assemble quality and correlation with GC-content, min mappability and genome size. (a) Distribution of TELL-Seq assembly quality indicators under different sequencing depth for clinical and standard strains. (b) Correlation between assemble quality indicators and GC-content, min mappability and genome size.

## Discussion

In our study, we for the first time showed that TELL-Seq produces near-complete genome assemblies across a wide range of bacterial pathogens, comparable to PacBio but with only 1/6 of cost and 1/15000 of DNA quantity. With the Tell-Link *de novo* assembly workflow, 200× depth of coverage guaranteed a robust assembly performance, regardless of genome complexity. For most bacterial genomes of moderate complexity, a depth of 100× can be sufficient if cost is a concern. However, the assembly quality started to decrease when coverage was beyond 200×, possibly due to the limited ability of the assembly algorithm in handling too large of a dataset. Genome complexity, indicated by mappability, is crucial for assembly quality, not GC-content. Our conclusions were limited as only the official Tell-Link *de novo* assembly workflow was used in the experiments. A more comprehensive benchmarking would be beneficial to access differences amongst assembly algorithms. Our dataset would help develop better algorithms across different bacterial species. In conclusion, TELL-Seq is a cost effective and efficient approach for bacteria genomics, opening up unprecedented possibilities for large-scale *de novo* assembly analysis.

## Methods

### Sampling, DNA Extraction and Mappability assessment

12 Standard bacterial species with differing GC-content were obtained from the American Type Culture Collection (ATCC), while 12 corresponding clinical strains were obtained from Center for the Preservation of Microbial and Viral Strains of Zhejiang Provincial Center for Disease Control and Prevention. Genomic DNA was extracted using a genomic DNA extraction kit (QIAmp DNA Kit, QIAGEN, Germany), according to the manufacturer’s instructions. Mappability, a measure of how unique or repetitive regions in the genome are[15], was used as a proxy for assessing genome complexity. It was calculated for each reference using GenMap (v1.3.0) [15] with option -E 2 -K 30. Min and mean mappability were calculated across the whole genome.

### TELL-Seq library preparation and sequencing

TELL-seq libraries were constructed using a TELL-Seq Microbial Library Prep Kit (Universal Sequencing Technology). 0.1 ng of genomic DNA from each isolate were used for library construction. The TELL-seq libraries were quantified by Qubit dsDNA BR Assay kit (Thermo Fisher Scientific) and pooled for sequencing on Illumina NovaSeq 6000 with 2 × 150 paired-end reads, 18-cycle Index 1 reads and 8-cycle Index 2 reads based on the manufacturer’s protocols. TELL-Seq sequenced raw data were first processed by the Tell-Read v1.1.1u2 for barcode correction and filtering before proceeding to downstream analysis (https://universalsequencing.com/en-cn/pages/tell-seq-software).

### PacBio library preparation and HiFi sequencing

High-quality DNA samples were selected with major band >30kb, and randomly fragmented into 15-18kb length using a Covaris ultrasonic disruptor. The large fragment DNA was enriched and purified using magnetic beads. Damage repair and end repair were performed on the fragmented DNA. Stem-loop sequencing adapters at both ends of the DNA fragments were connected, and the failed fragments were remove using exonuclease. The constructed library was sequenced using the PacBio Sequel II/PacBio Sequel IIe platform. PacBio original sequencing reads are dumbbell-shaped sequences with both ends attached to the adapter, known as Polymerase reads. The raw data is broken down from the adapter and filtered out to obtain subreads, which are filtered according to the standard Filtering subreads by minimum length = 50. Based on subreads, ccs software is used to generate high-precision HiFi reads. Here, we set the criteria as min-passes=3, min-rq=0.99, which means that all read quality values are above Q20. This part of data is an effective input for downstream analysis.

### *De novo* assembly and quality assessment

Tell-Link v1.1.1u2 (https://universalsequencing.com/en-cn/pages/tell-seq-software) utilizes the linked-read data derived from the Tell-Read output to construct a barcode-aware assembly graph, assemble contigs, and perform scaffolding. When TELL-Seq reads were considered as short reads, adapters were trimmed and quality checked using fastp (v.0.23.4) [16] with default option. The clean reads were then assembled using the Unicycler (v0.5.0) [17] pipeline with default option. PacBio HiFi reads were first trimmed with Filtlong (v0.2.1) (https://github.com/rrwick/Filtlong) with options --min_length 1000 --keep_percent 95. And then assembled with Flye (v2.9.3-b1797) with option --meta --pacbio-hifi.

To assess assembly quality, Illumina short read assembly and TELL-Seq assembly were benchmarked against PacBio assembly for clinical strains and against reference for standard strians. QUAST (v5.0.2) [18] was employed to calculate evaluation metrics, while MUMmer (v3.1) [19] was utilized for the analysis of collinearities plots.

### Downsampling analysis for TELL-Seq genome assembly

Raw TELL-Seq reads were downsampled to 50×, 100×, 150×, 200×, and 250× depths with seqtk (https://github.com/lh3/seqtk). Same assessment strategies were undertaken as above for assembly quality. Statsmodels and scipy libraries (v1.13.1) in Python (v3.9) was used to calculate the Spearman correlation coefficients, the corresponding p-values and adjusted p-values between assembly quality statistics and quality control indicators or genomic characteristics (mappability, genome size or GC-content).

## Supporting information

all supplemental tables

## Funding Information

This work was funded by the Key R&D Program of Zhejiang Province (2024C03216) to ZJCDC and the Ministry of Science and Technology, China (2020YFC2008801) to AIIT, PKU.

## Acknowledgement

We would like to thank Mr. Longfei Xie for his assistance with coordinating and conducting this study.

## Contribution

BW, JJ and XL conceived and designed the experiments; ZW, ZY did the experiment; TZ, LS, SL and XL performed the data analysis; TZ, SL, LS, YY, XC, BL, SZ and XL developed the data analysis workflows; ZW, TZ, SL, LS, BW, JJ and XL wrote the manuscript; XZ helped managing and coordinating the research project. All authors contributed to the revision and review of the manuscript.

## Conflict of Interests

No conflict of interests.

## Supplemental Material

**Fig S1.**
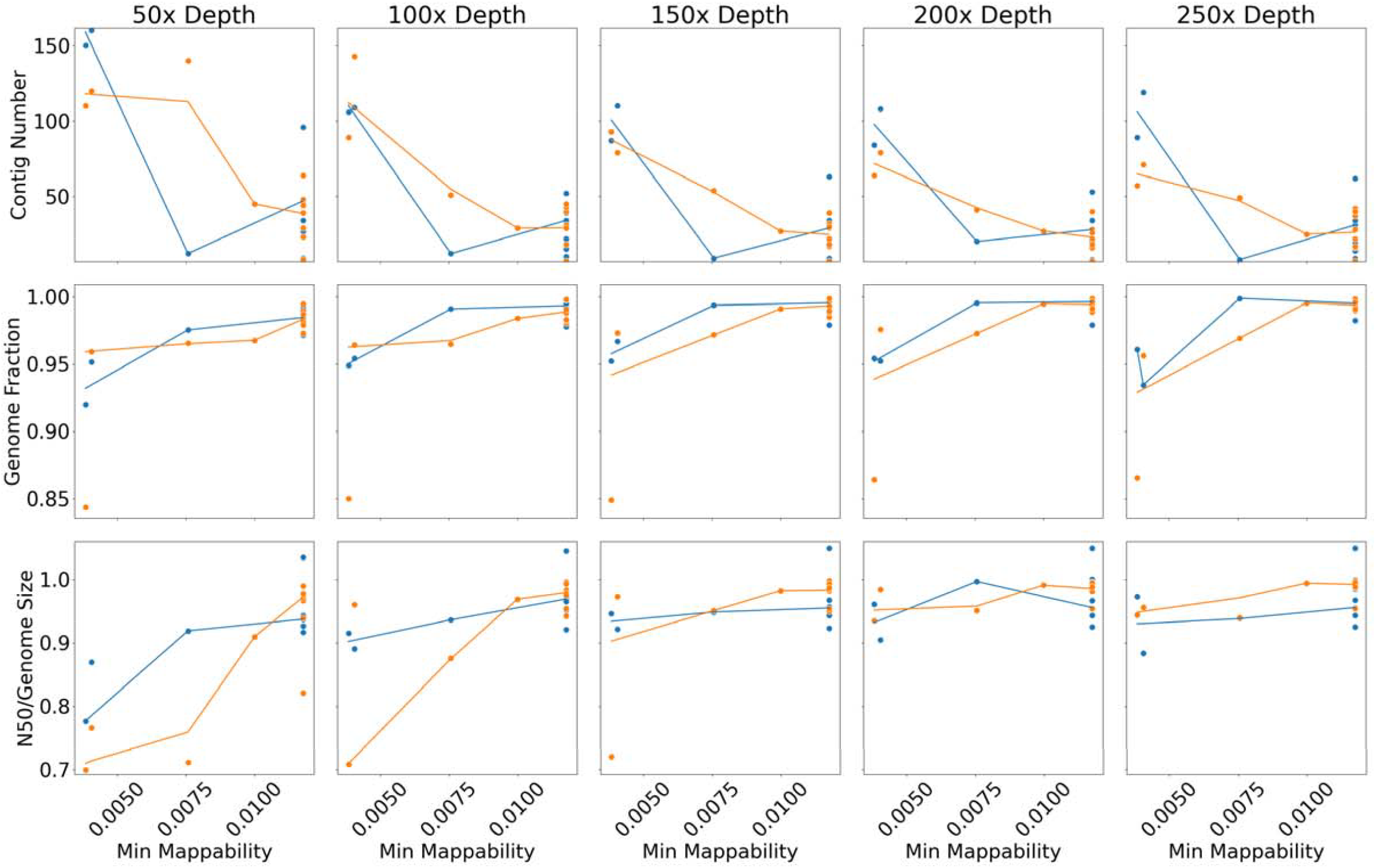
Scatter Plot of Minimum Mappability *vs*. Assembly Quality Across Different Sequencing Depths. Scatter plot showing the relationship between minimum mappability and assembly quality across different sequencing depths.

**Fig S2.**
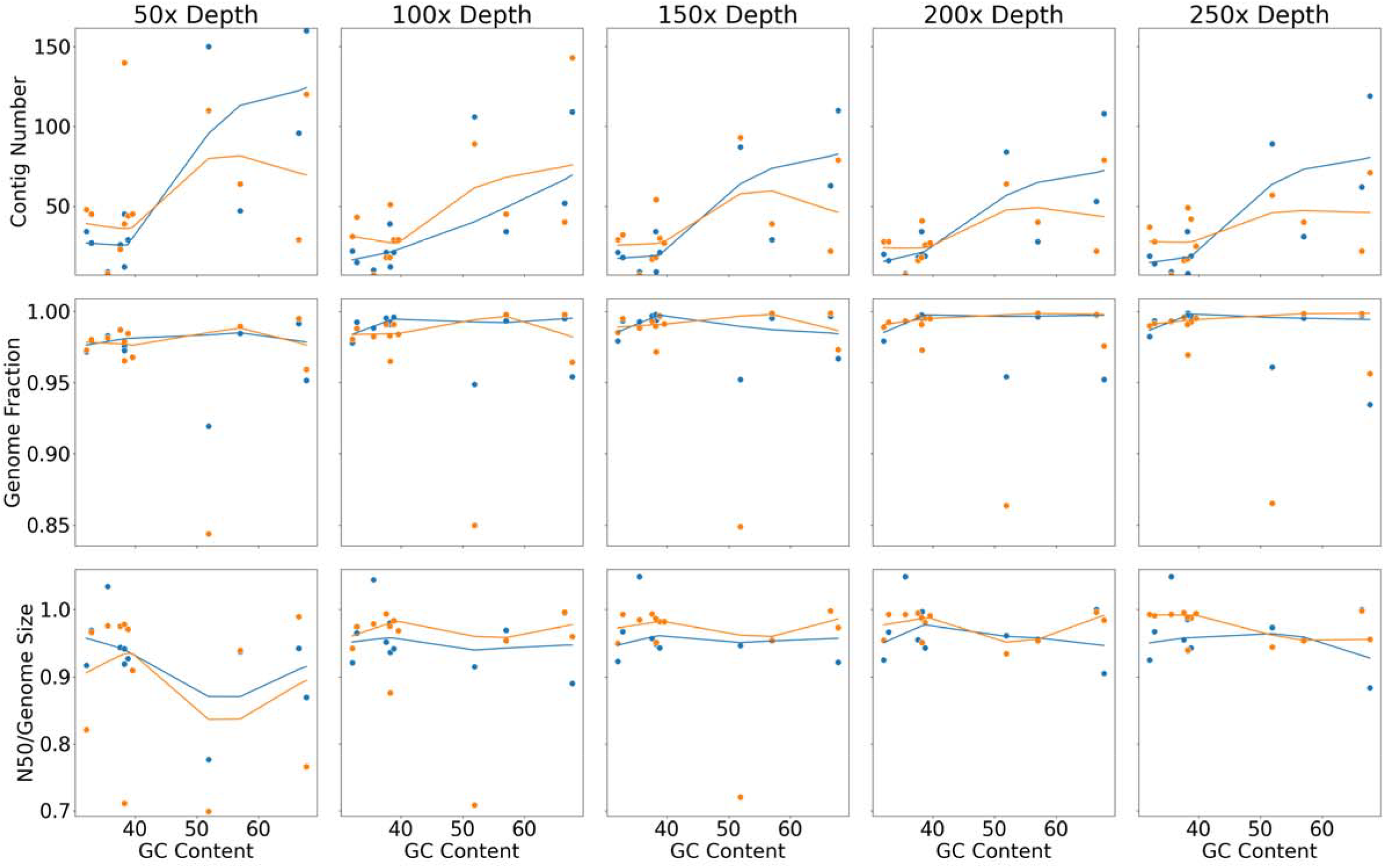
Scatter Plot of GC-content *vs*. Assembly Quality Across Different Sequencing Depths. Scatter plot showing the relationship between GC-content and assembly quality across different sequencing depths.

**Fig S3.**
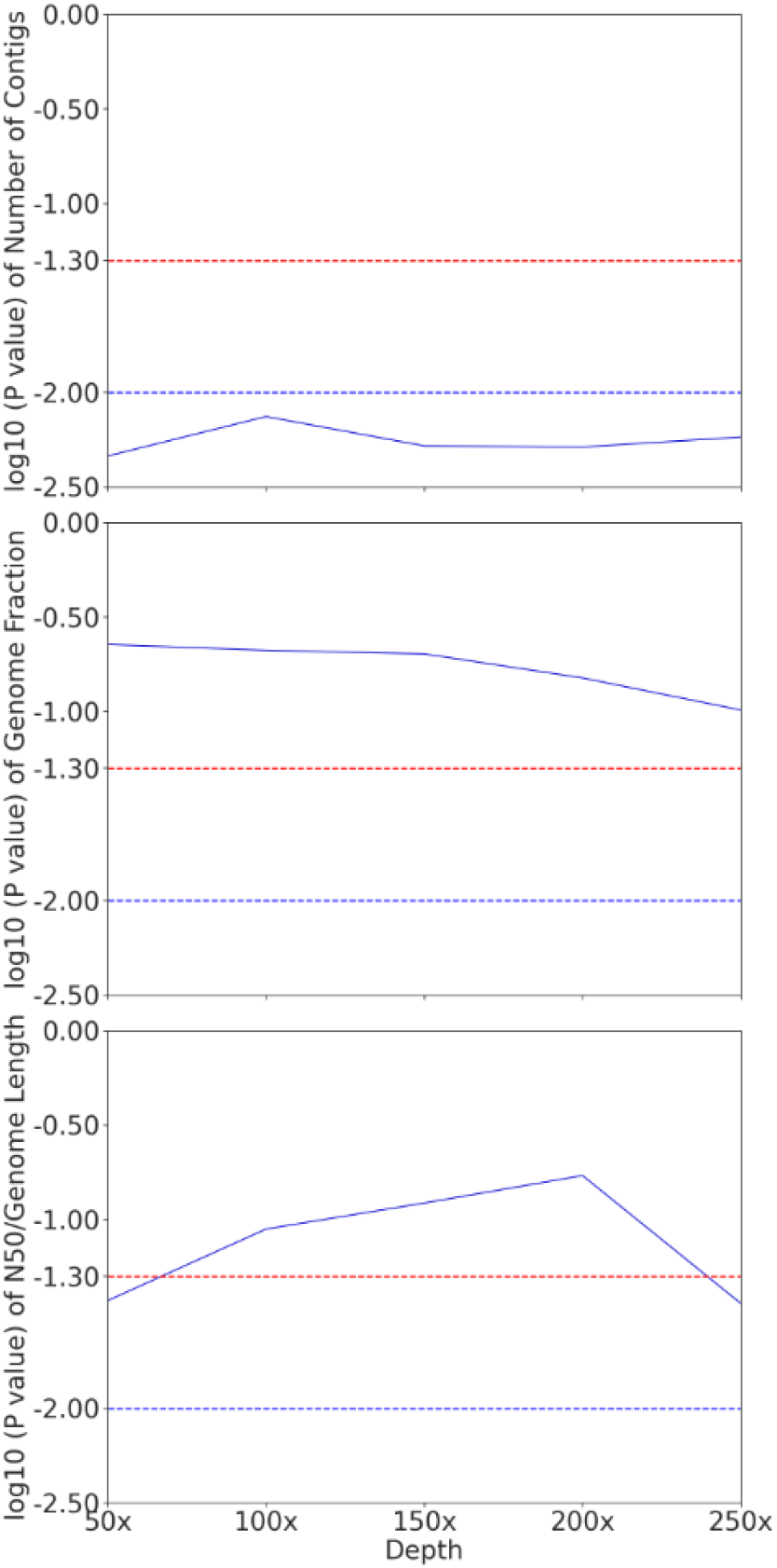
Wilcoxon Sum Rank test of assembly quality statistics between Gram positive and negative across different sequencing depths.

**Tab S1: Overview of sequenced bacteria genomes**

**Tab S2: Summary of PacBio raw sequencing data quality of clinical samples**

**Tab S3: Summary of PacBio assembly quality of clinical samples**

**Tab S4: Summary of TELL-Seq data quality**

**Tab S5: BioSample Objects**

## Notes

### Competing Interest Statement

The authors have declared no competing interest.

